# Macronutrients modulate resistance to infection and immunity in Drosophila

**DOI:** 10.1101/498493

**Authors:** Fleur Ponton, Juliano Morimoto, Katie Robinson, Sheemal S Kumar, Sheena Cotter, Kenneth Wilson, Stephen J Simpson

## Abstract

- Immunity and nutrition are two essential modulators of individual fitness
- However, while the implications of immune function and nutrition on an individual’s lifespan and reproduction are known, the interplay between feeding behaviour, infection, and immune function, remains poorly understood.
- In this study, we used the fruit fly, *Drosophila melanogaster*, to investigate how infection through septic injury modulates nutritional intake, and how macronutrient balance affects survival to infection by the pathogenic Gram-positive bacterium *Micrococcus luteus*.
- Our results show that infected flies maintain carbohydrate intake, but reduce protein intake, thereby shifting from a protein-to-carbohydrate (P:C) ratio of ~1:4 to ~1:10 relative to non-infected and sham-infected flies.
- Strikingly, we found that the proportion of flies dying after *M. luteus* infection was significantly lower when flies were fed a low-P high-C diet, revealing that flies shift their macronutrient intake as means of nutritional self-medication against bacterial infection.
- This is likely due to the effects of macronutrient balance on the regulation of the constitutive expression of innate immune genes, as a low-P high-C diet was linked to an up-regulation in the expression of key antimicrobial peptides.
- Together, our results reveal the intricate relationship between macronutrient intake and resistance to infection, and integrate the molecular cross-talk between metabolic and immune pathways into the framework of nutritional immunology.

## Introduction

Infection and nutrition are intricately and intimately linked (Kelley and Bendich, 1996; Sheldon and Verhulst, 1996; Samartin and Chandra, 2000; Rolff and Siva-Jothy, 2003; Cunningham-Rundles et al., 2005; Bauer et al., 2006; Calder, 2006; Falagas and Kompoti, 2006; Amar et al., 2007; Klasing, 2007; Wu et al., 2007; Ayres and Schneider, 2009; Falagas et al., 2009; Lazzaro and Little, 2009; Sorci and Faivre, 2009; Hawley and Altizer, 2010; Ponton et al., 2011a; Schmid-Hempel, 2011; Huttunen and Syrjanen, 2013; Ponton et al., 2013; Genoni et al., 2014; Martinez et al., 2014, Vogelweith et al. 2015,). Recent studies have allowed a detailed molecular understanding of the cross-regulation between nutrition and immunity, with nutrient sensing pathways being identified as important regulators of innate immunity (Becker et al., 2010; Martin et al., 2012; Varma et al., 2014). Immunity can be activated independently to an infection and this regulation can act under conditions of fluctuating nutrient availability. While the underlying mechanisms are far from being fully understood, the relationship between diet, diet-induced metabolic diseases and infections is clearly multi-factorial, with impairments of immune function playing a key role (Martí et al., 2001; Nave et al., 2011). Better understanding the nutritional components that influence immunity and resistance to infection is an important challenge with implications for animal and human health.

There is an ongoing debate on the effects of diet on immune responses to infections. Food deprivation, and/or protein shortage has been reported to negatively affect immunity responses and survival after infection (Siva-Jothy and Thompson, 2002; Pletcher, Macdonald, Marguerie et al., 2002, Brunner et al., 2014) with infected hosts selecting a protein-biased diet that provided them with a better survival after infection (Lee et al., 2006; Povey et al., 2009; Povey et al., 2014). In *Drosophila*, while diet restriction has been shown to decrease the capacity of the host to clear the infection (i.e., “resistance”), it provided the host with the ability to reduce the damage of the infection on its health, also called “tolerance” (Ayres and Schneider, 2009, 2012). More recently, it has been shown that yeast restriction affects tolerance specifically to one strain of bacterium in a time-dependent manner; however, no effect on resistance was detected (Kutzer and Armitage, 2016, see also Miller and Cotter, 2017 and Howick and Lazzaro, 2014).

Finally, a negative effect of protein and/or a positive effect of carbohydrate on resistance have been revealed (Graham et al., 2014; Kay et al., 2014; Mason et al., 2014) with, for instance, female *Drosophila* fed an holidic diet supplemented with glucose having greater survival following infection with the gut pathogen *Vibrio cholarae* (Galenza et al., 2016). Although there is a clear effect of diet composition on resistance to infection and immune state, dietary manipulations have usually focused on changing single nutrients or varying the caloric content and nutrient ratio simultaneously, which hinders the ability to specifically measure the effects of food components and/or caloric content on immunity [but see (Cotter et al., 2011)]. There is now evidence that considering the interactive effects of nutrients is essential and offers a more ecologically relevant understanding (Cotter et al., 2011; Simpson and Raubenheimer, 2012; Simpson et al., 2015).

Here, we explored the nutritional responses of *Drosophila melanogaster* after bacterial challenge and the consequences of such responses for survival following infection. We performed a detailed investigation of the dietary modulation of constitutive innate immune gene expression in an age-dependent manner. The effects of nutrition were measured through a geometric manipulation of the dietary protein and carbohydrate balance. Our observations unveiled nutritional regulations of innate immune gene expression and resistance to bacterial infections, and link these findings to nutritional self-medication.

## Results

### Bacterial infection induces a shift in dietary choice to a low P:C diet

We first hypothesized that infection through septic injury with the pathogen *Micrococcus luteus* would modulate the nutritional selection of *Drosophila melanogaster*. Adult flies were offered a choice between two capillaries filled with either a sucrose or a yeast solution, and food intake was measured every two days for six days (Ja et al., 2007) (see Methods). While non-, sham- and *M. luteus*-infected flies ingested similar quantities of carbohydrate (cumulative consumption of carbohydrate over six days, One-way ANOVA, F_2,36_=1.775, p=0.185, Supplementary Table 1), the consumption of protein was significantly different between the treatments (i.e., cumulative consumption of protein for six days, One way ANOVA, F_2,36_=5.853, p=0.007, Supplementary Table 1). Protein consumption was the lowest for flies infected with *M. luteus* and was the greatest for sham-infected and non-infected flies (Fig. 1). This reduction in protein intake by infected flies resulted in a marked change in the ingested dietary P:C ratio, such that flies infected with *M. luteus* balanced their diet to a P:C ratio close to 1:9.6 (i.e., 9% protein, Fig. 1) and non- and sham-infected to a P:C ratio of 1:3.8 (i.e., 20% protein) and 1:3.2 (i.e., 25% protein), respectively (Fig. 1).

**Figure 1.**
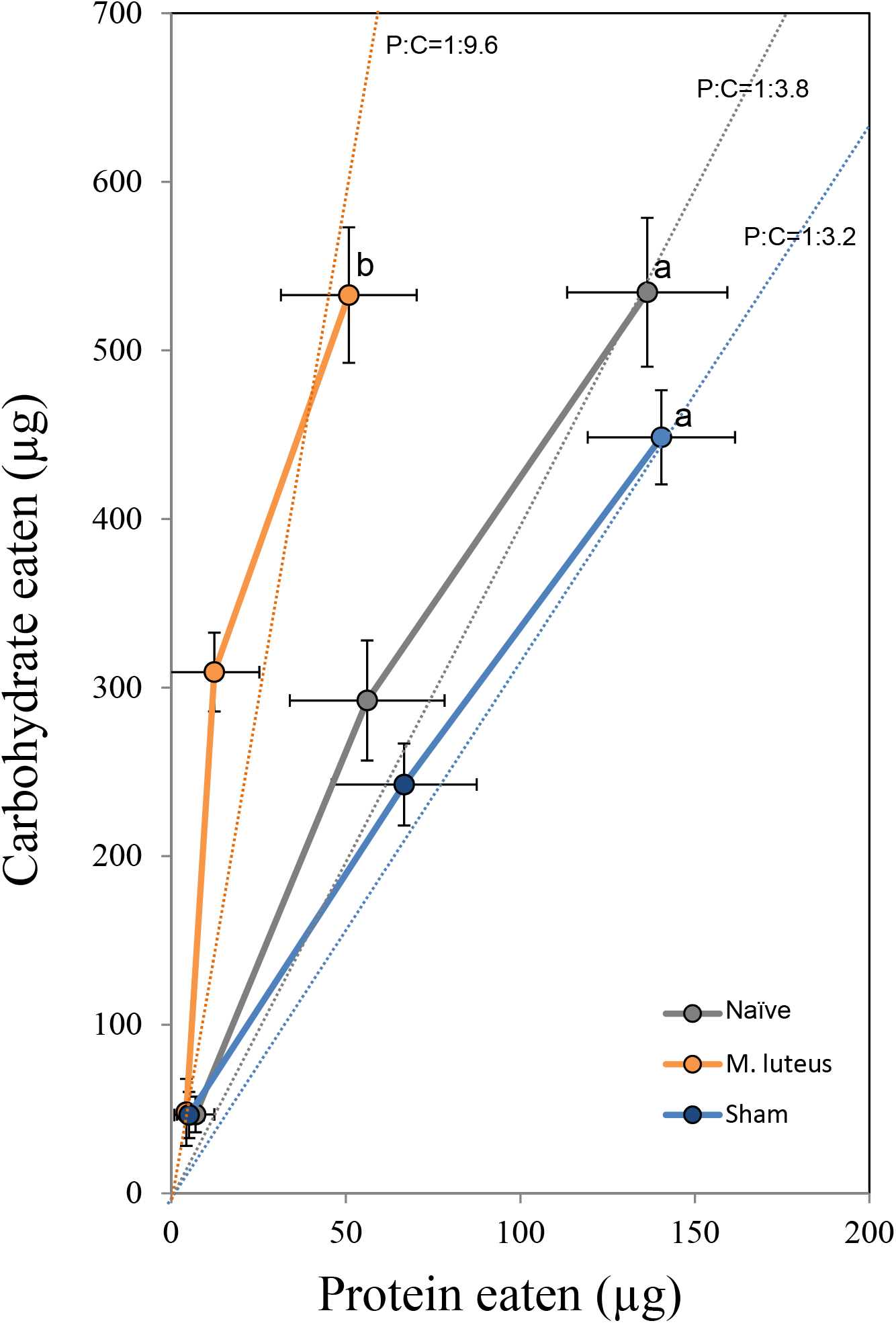
Cumulative protein–carbohydrate intake (mean+s.e.m.) trajectories at two-day intervals over six days. Dotted lines represent protein-to-carbohydrate ratios (P:C). Letters indicate significant HSD Tukey’s pairwise comparisons (p ≤ 0.05).

The level of expression of several immune genes was measured for the different treatments six hours post-challenge (see Methods). Infection with *M. luteus* induced the greatest changes in immune gene expression (Fig. 2). Relative to naïve flies, infection by *M. luteus* triggered the enhanced expression of all of the antimicrobial peptides assayed (i.e., AttaA, CecA, CecC, DipB, Def, Mtk, Supplementary Table 2), as well as molecules involved in the transduction of the immune signal (i.e., spz and Dif, Supplementary Table 2). However, no significant effect of infection was detected on the level of expression of the two receptors involved in the recognition of pathogens that we assayed (i.e., GNBP2 and PGRPSA, Supplementary Table 2 and Fig. 2). Even though the expression levels of spz, CecA, CecC and DiptB were significantly greater in sham-infected flies compared to non-infected insects, levels of expression of these genes remained more elevated in infected individuals compared to sham-infected and non-infected individuals (Fig. 2).

**Figure 2.**
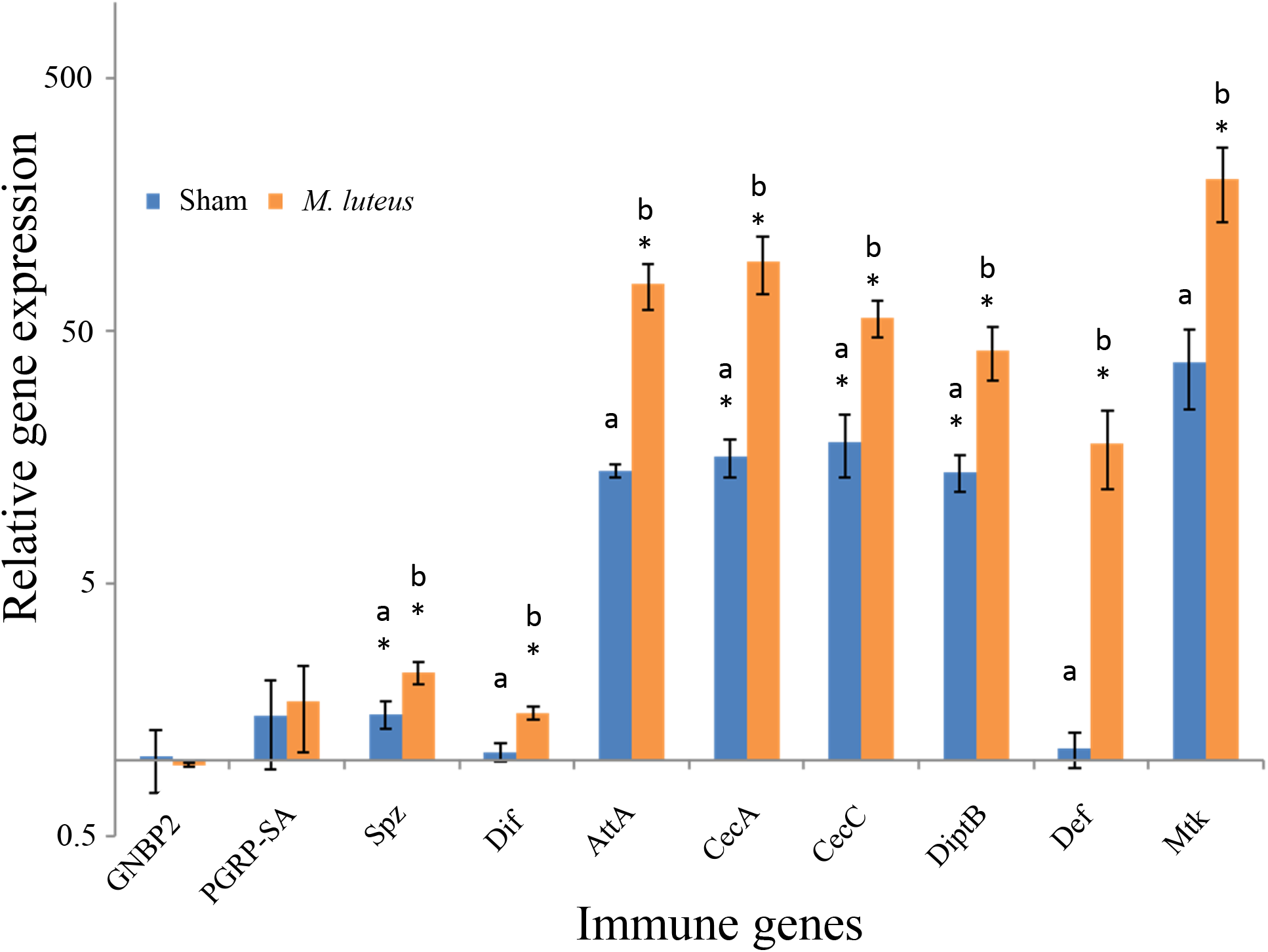
Relative immune gene expression (relative to naïve flies, mean+s.e.m.). Letters indicate significant HSD Tukey’s pairwise comparisons (p≤0.05) between sham-injured and *M. luteus*- infected flies, stars (*) indicate significant Bonferroni pairwise comparisons (p≤0.05) against naïve flies.

These first results show that when flies are infected with *M. luteus*, they shift their nutritional choice to a carbohydrate-biased (lower P:C) diet, which is above and beyond the stress of physical injury (i.e. compare sham-infected vs. infected).

### A low P:C diet can improve survival post-infection

We then hypothesized that the shift to a low P:C diet observed for infected flies had survival significance. In this second experiment, non-, sham- and *M. luteus*-infected flies were fed one of three diets (high, medium and low P:C in a no-choice experiment) and survival was followed. As expected, the interaction between the dietary P:C and the treatment significantly influenced survival rate of flies (Cox regression, Treatment X Diet: χ^2^=26.97, df=4, p<0.001, Treatment: χ^2^=66.28, df=2, p<0.001, Diet: χ^2^=606.57, df=2, p<0.001). Survival was reduced on higher P:C diets for the three groups of flies compared to the two other diets (Fig. 3). However, while naïve flies survived in similar proportions on medium and low P:C diets (i.e., 24% and 4% protein) (Log Rank pairwise comparisons, p>0.05, Fig. 3A), *M. luteus*- and sham-infected flies survived significantly better on the low P:C diet (i.e., 4% protein) compared to the medium P:C diet (i.e., 24% protein diet) (Log Rank pairwise comparisons, p≤0.05; Fig. 3B&C). At day 15, the interaction between diet and treatment significantly influence the percentage of dead flies (Supplementary Table 3A). Flies on high P:C diet had great percentage of death (Supplementary Table 3B; post hoc p>0.05). On medium P:C diet, we found a greater percentage of dead flies for flies infected with *M. luteus* compared to sham-infected and naïve treatments (Supplementary Table 3A; post hoc test, p≤0.05). Percentage of dead flies was also greater for sham-infected flies compared to naïve individuals. On low P:C diet, however, the percentage of dead flies was not significantly different between the 3 treatments (Supplementary Table 3A; post hoc test, p>0.05).

These results suggest that flies better resisted infection by *M. luteus* when fed a low-protein, high-carbohydrate diet.

**Figure 3.**
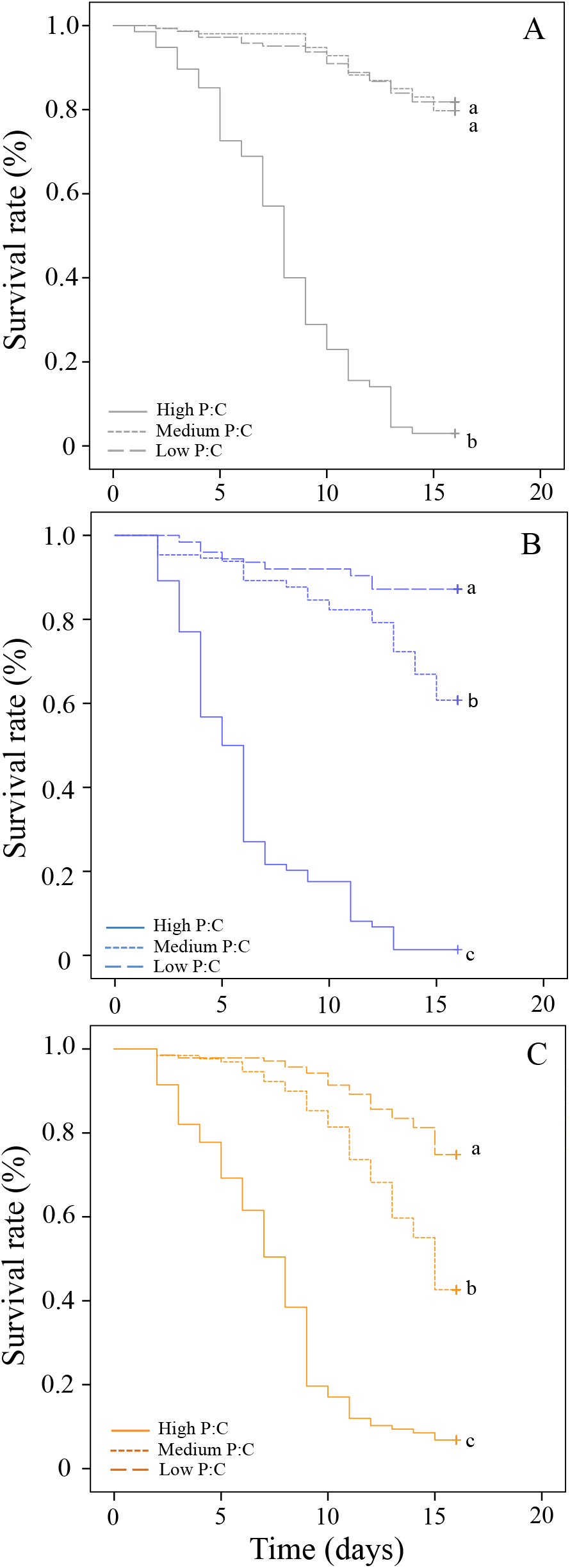
Survival curves of flies fed three diets varying in the protein-to-carbohydrate ratio (P:C) [i.e, P:C=1:1 (high), 1:4 (medium), and 1:32 (low)] after treatment (Naïve, Sham- and *M. luteus-infected*). Letters indicate significant pairwise comparisons (p ≤ 0.05).

### The P:C ratio influences the constitutive expression of antimicrobials

We next investigated the underlying mechanisms mediating the effect of carbohydrate-biased diet on immune state. Given our findings, we hypothesized that a low-protein, high-carbohydrate diet stimulates the expression of immune genes. We measured the expression of 21 genes involved in the integrated response to pathogen infection, beginning with pathogen recognition receptors, transduction of the immune signals, and antimicrobial peptides (AMPs) for flies fed seven isocaloric diets varying in the P:C ratio (the percentage of dietary protein was used as a descriptor in the analyses and figures, see Methods and Supplementary Table 4). Flies were sampled at three key points on their life expectancy curves (i.e., 25, 50, 75% mortality, see Supplementary Fig. 1 and Supplementary Table 5).

Our data show that immune genes expression levels were significantly influenced by the ratio of protein-to-carbohydrate in the diet (expression data pooled for all AMPs; Kruskal-Wallis test: 25% mortality, χ^2^=43.619, df=6, p≤0.001, N=157; 50% mortality, χ^2^=27.279, df=6, p≤0.001, N=158; 75% mortality, χ^2^=51.345, df=6, p≤0.001, N=153; Fig. 4). Expression level of the genes coding for AMPs was overall negatively associated with dietary P:C and this was observed at the three sampling times, though there is some suggestion of non-linear trends in the earlier sampling points. Expression of genes coding for immune receptors was significantly influenced by dietary P:C, however we did not detect any clear pattern of variation (Supplementary Fig. 2 and Supplementary Table 6). Diet composition did not influence expression levels of genes coding for molecules involved in the transduction of the immune signal (Supplementary Table 6).

**Figure 4.**
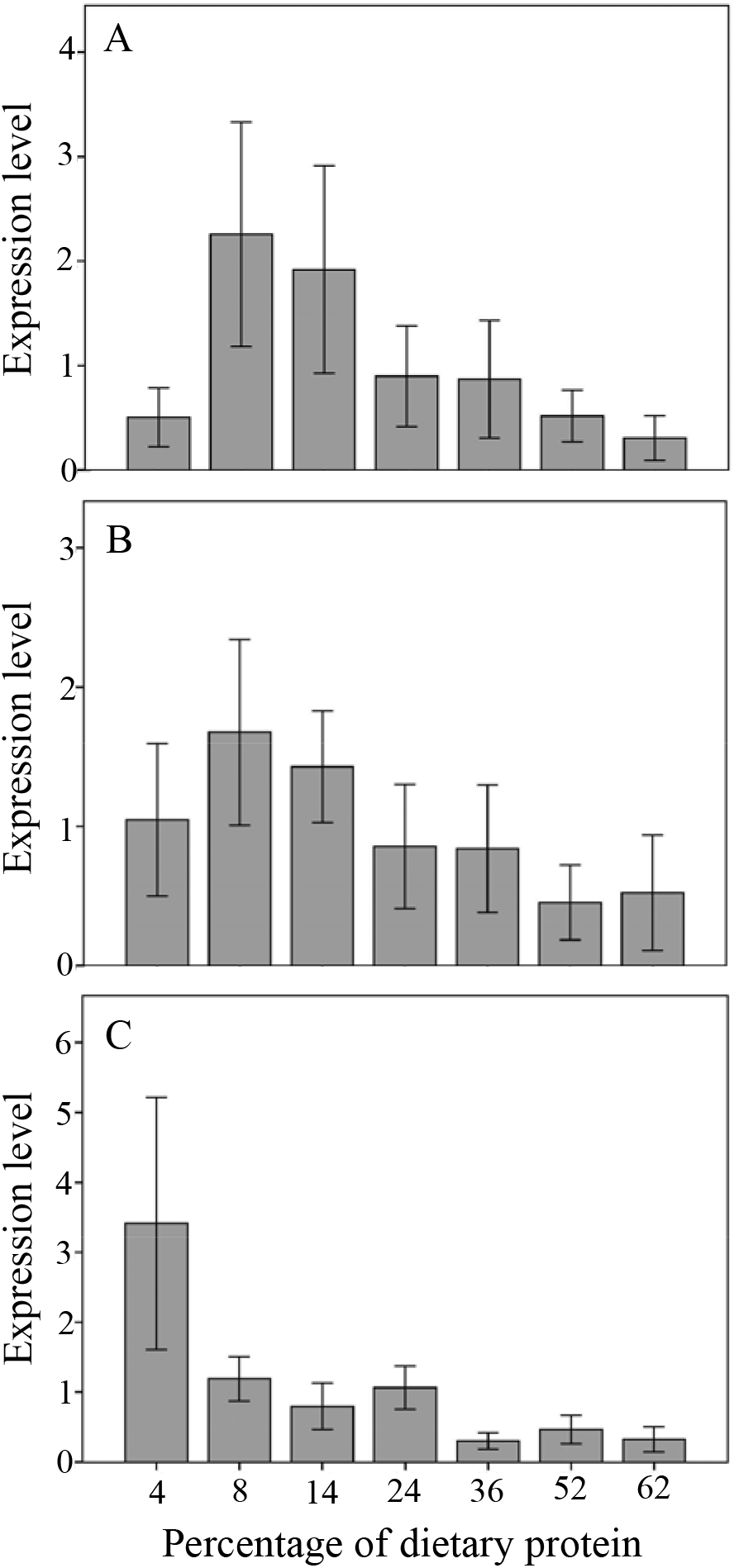
Expression levels of antimicrobial genes (mean±SE) according to the percentage of protein in the diet at: A. 25% mortality, B. 50% mortality, C. 75% mortality.

When we looked in more details at the effect of dietary P:C on the expression level of the specific genes, we found a significant negative non-linear relationship between the level of expression and the percentage of dietary protein for six out of nine genes coding for antimicrobial peptides (Fig. 5), which reveals that antimicrobial peptide expression is tightly linked with the macronutrient balance in the diet. This diet-dependent effect on antimicrobial peptide expression was consistent throughout the flies’ lifespan (see Supplementary Fig. 3).

**Figure 5.**
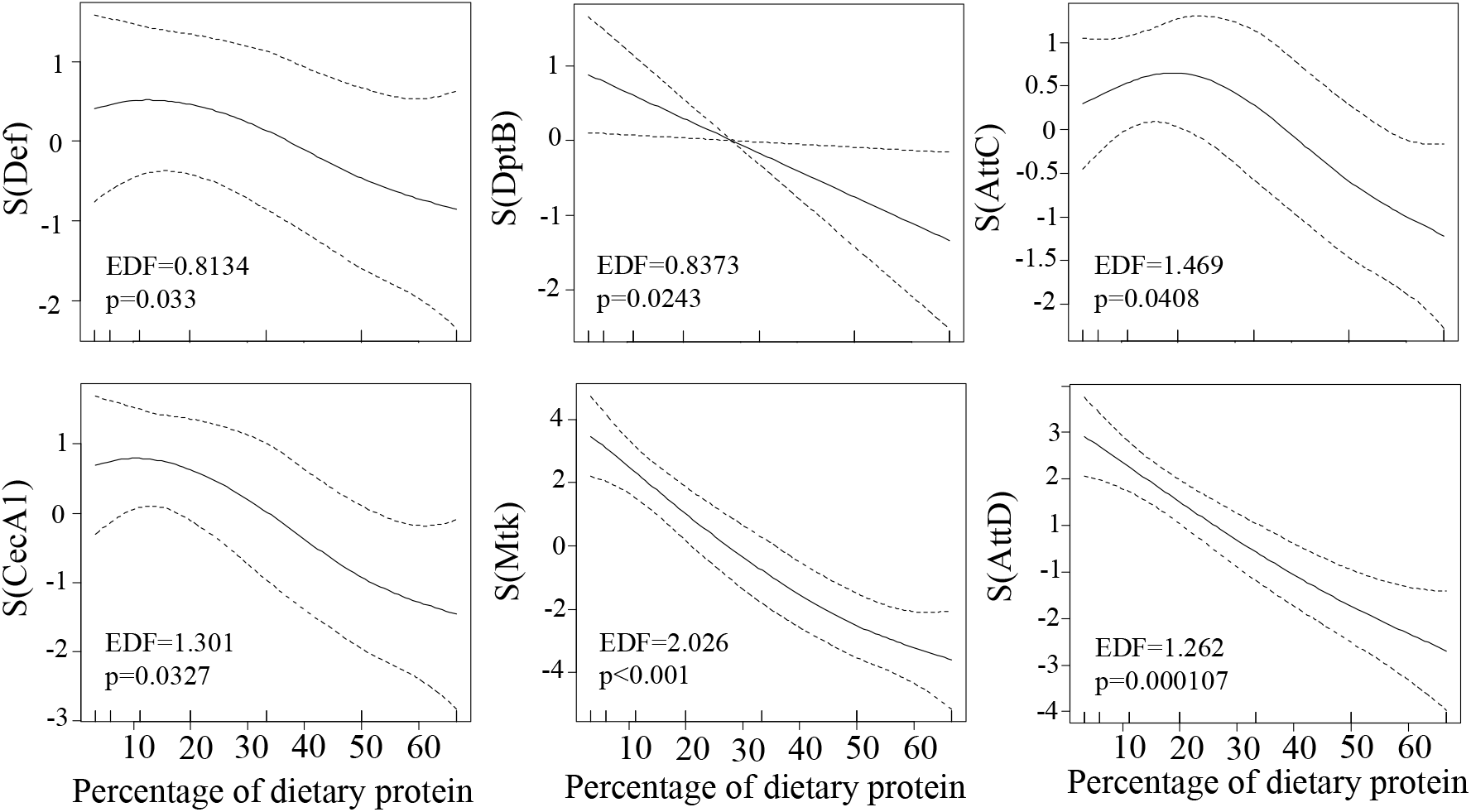
Estimated nonparametric smooths of antimicrobial gene expression levels from the generalized additive model according to the percentage of dietary protein at 50% survival (Def= defensin, DptB= DiptericinB, AttC= AttacinC, CecA1=CecropinA1, Mtk= Metchnikowin, AttD= AttacinD).

Interestingly, we found that the effect of dietary P:C can vary depending on the sampling point and the specific gene. For example, for the pattern recognition proteins, gene expression was positively associated with P:C for PGRPSC2, GNBP1 (at 25% mortality only for both genes) and PGRPLC (at 50% mortality only), whereas there was a negative association for PGRPSA (at 25% and 75% mortality) and PGRPSB1 (at all sampling points) (see Supplementary Fig. 3). Expression of genes coding for proteins involved in the immune-signal transduction (i.e., Dif, Imd, Relish, Thor, Toll, Spätzle) was generally not significantly influenced by dietary P:C (Supplementary Fig. 3). Together, these results suggest that a carbohydrate-biased diet can maintain a higher constitutive expression of antimicrobial peptides that might allow flies to better fight infections and injuries.

## Discussion

Our results confirm the key role of protein and carbohydrate in immunity and resistance to infection. Although the dietary ratio of protein to carbohydrate (P:C) modulated expression of genes linked to innate immunity, it did not affect all immune molecules in the same way (see also (Cotter et al., 2011)). While AMP expression levels were overall negatively affected by the relative amount of protein in the diet, the effects of dietary P:C on molecules involved in the recognition of pathogens depended on gene identity. In addition, overall, no effect of dietary P:C was detected on molecules involved in the transduction of the immune signal.

In a diet-choice experiment, we showed that flies infected with *M. luteus* decreased their protein consumption while maintaining carbohydrate intake at the same level as non-infected individuals. While anorexia – i.e. a sharply decreased overall food intake – has been proposed to enhance tolerance and/or resistance [see for instance (Ayres and Schneider, 2009; Adamo et al., 2010)], our results strongly support the notion that modulation of macronutrient balance, rather than overall calories, may underpin this effect (Cotter et al., 2011) [see also (Fontana and Partridge, 2015) for a similar discussion on the effects of nutrients on longevity]. Furthermore, while sham-infected individuals survived better on low P:C, we did not observe any significant shift in their diet choice. The magnitude of the nutritional effects in sham-infected flies was a bit smaller compared to infected flies and might explain these results. More investigations are needed to fully understand this result.

The cross-regulation between immune and metabolic pathways involves molecules such as Forkhead box O (FOXO), target of rapamycin (TOR) and 5’ AMP-activated protein kinase (AMPK) in mammals and *Drosophila* (Becker et al., 2010; Martin et al., 2012; Seiler et al., 2013; Abdel-Nour et al., 2014). Inhibition of TOR signaling has been shown to promote a pro-autophagic and inflammatory environment that is essential for clearing infections (Chakrabarti et al., 2012; Martin et al., 2012), which might result from nutrient deficiencies, such as amino-acid deprivation following host membrane damage (Tattoli et al., 2012). Varma et al. (2014) have also shown that inhibiting the TOR pathway using mutants and the drug rapamycin results in an enhanced expression of several AMPs in *Drosophila* (Varma et al., 2014). Interestingly, this system can be manipulated by pathogens that have evolved ways to maintain TOR complex activity in an amino acid-independent manner (Clippinger et al., 2011). Dietary protein to carbohydrate ratio was predicted to modulate TOR activation, as shown recently in mice (Simpson and Raubenheimer, 2009). As a result, we predict that antimicrobial peptides are up-regulated on high-carbohydrate, low-protein diets because of low TOR activation [see also (Varma et al., 2014)]. However, we did not detect a prophylactic effect of carbohydrate per se, as it has been suggested in an earlier study (Galenza et al., 2016).

Micronutrients are also important food components that can modulate immunity (Calder, 2017 for more details). We here approached foods as mixtures of macronutrients (and correlated content of micronutrients) and do not specifically address the effects of micronutrients on fly immunity. More investigations through specific manipulations of dietary micronutrient content would allow to further explore the role of micronutrients on immunity and resistance to infection in drosophila.

Our results show that flies better survived infection by *M. luteus* through modulating their macronutrient intakes 15 days after infection. Self-medication has been traditionally seen as animals using molecules such as secondary plant compounds or other non-nutritive substances with antiparasitic activity (Raubenheimer and Simpson, 2009; de Roode et al., 2013). However, our work reinforces the idea that self-medication can happen through modulating macronutrient selection to stimulate the immune response and potentially compensate for the negative effects of the infection on fitness traits [see for instance (Abbott, 2014; Povey et al., 2014; Galenza et al., 2016; Bashir-Tanoli et al. 2014)]. Specific appetites for protein and carbohydrate mediate diet selection and food intake (Simpson et al., 2015) and better understanding the links and relationships between macronutrients, the immune state, and resistance to infection will inform the use of nutritional interventions as tools for improving the health of animal and human populations.

## Material and methods

### Experimental infection

One day-old adult female flies (Canton-S, stock from Bloomington) were experimentally infected using a solution of freshly grown *Micrococcus luteus* (ATCC 10240) at OD600=0.5. Flies were anaesthetized under CO2 and pricked in the thorax using a dissecting pin that was beforehand dipped in the bacterial solution [see (Apidianakis and Rahme, 2009)]. We also generated sham-infected flies using a pin dipped in ethanol (70%). As negative controls, we used non-infected, non-injured flies (i.e., naïve flies). Flies were left to recover from pricking for half an hour. Survival immediately after the infection was ~95%.

### Nutritional intake target

Infected, sham-infected, and naïve flies were individually provided with two 5μl microcapillary tubes (Drummond Microcaps) filled with liquid diets (n=20 flies per treatment at the start of the experiment): one diet consisted of autolyzed yeast (MP Biomedicals, catalogue no. 103304) at 180g/L and the other, of sucrose at 180g/L. The two solutions were prepared in sterile, distilled water. Intake was measured against a scale bar by height difference in the column of liquid within the microcapillary every 2 days for 6 days [see (Lee et al., 2008; Ponton et al., 2015)]. Total quantities of protein and carbohydrate ingested were compared using One-way ANOVA type II and post-hoc tests (Tukey’s HSD).

### Immune gene expression levels using RT-qPCR

We investigated the expression of immune genes of the IMD and Toll pathways using reverse transcription quantitative PCR (RT-qPCR). We used 1d-old adult female infected, sham-infected, or control. After pricking, flies were left to recover for half an hour before being transferred to P:C=1:4 (3 replicate cages per treatment). After 6h, flies were dissected (i.e., eggs removed), preserved in RNA later (Ambion) and stored at −80 °C. RNA was extracted for 10 to 15 flies per cage (see below for more details on RNA extraction). Complementary DNA was generated using the QuantiTect Reverse Transcription Kit (Qiagen). Triplicate cDNA aliquots for each sample served as templates for quantitative PCR using SYBR Green PCR Master Mix (Applied Biosystems). Amplification reactions were performed in 10μl total volumes with 4.5μl of cDNA (diluted 1:90) and 100 to 200 nM of each primer [see (Ponton et al., 2011b) for the primer sequences of reference genes Rpl32 (Ribosomal protein l32, CG7939) and Ef1 (Elongation factor 1, CG1873); see Supplementary Table 6 for the primer sequences of target genes], in 384-well optical plates under the following sequential conditions: incubation at 50°C for 2 min, 95°C for 10 min, followed by 45 cycles of 95°C for 15 s and 60°C for 1 min. RT-qPCR efficiency was determined for each gene and each treatment using second derivative method. Relative standard curves for the gene transcripts were generated with serial (5x) dilutions of cDNA (i.e. 1/20, 1/40, 1/80, 1/160 and 1/320). Stock cDNA used for the relative standard curves consisted of a pool of cDNA from the different samples. No template and no RT controls were run for each primer pair. Target gene expression levels were normalized by reference gene expression levels. Expression levels were given relative to the control treatment (i.e., non-injured, non-infected flies) for each gene and compared between treatments using generalized linear model analysis (GLM) followed by post-hoc tests (Tukey’s HSD). Effect of the percentage of dietary protein on each single gene expression level was analyzed using generalized additive model analysis (GAM).

### Effect of dietary manipulation on resistance to infection

One day-old adult female flies were infected as described above. Flies were left to recover for half an hour before being transferred to experimental cages, and split into groups of 50 individuals fed with three solid diets varying in the P:C ratio. Foods varied in hydrolysed yeast (Y) and sucrose (S) content. The Y:S concentration was 180g/1. Macronutrient compositions were calculated based on autolyzed yeast [MP Biomedicals, catalogue no. 103304 containing 62% protein]. Each diet contained 0.01% phosphoric acid and 0.1% proprionic acid as antimould agents and were prepared in sterile, distilled water. Dietary treatments were defined as “high P:C ratio” (i.e., P:C=1:1 or 52% P), “medium P:C ratio” (i.e., P:C= 1:4 or 24% P) and “low P:C ratio” (i.e., P:C=1:32 or 4% P). Three replicates cages for *M. luteus-* and sham-infected flies and two replicate cages with naïve flies were run in parallel for each dietary treatment. Lifespan was followed for 16 days with dead flies counted daily. Flies that died from 0 to 6h post-infection were removed from the analyses since we could not assess if the death was directly caused by the infection or the dietary treatment. Kaplan-Meier lifespan curves were analyzed using Cox regression and Log Rank Mantel-Cox tests.

### Immune gene expression levels using Taqman Low-Density Array (TLDA) cards

#### Dietary treatments

Foods varied in hydrolysed yeast (Y) and sucrose (S) content. The seven Y:S ratios used were 1:14, 1:7, 1:3.5, 1:1.6, 1:0.7, 1:0.2, or 1:0; yielding protein-to-carbohydrate ratios of 1:21, 1:11, 1:5, 1:2.5, 1:1, 3:1 and 1:0, respectively; and percentages of protein (w/w(Y+S)) of 4%, 8%, 14%, 24%, 36%, 52% and 62%, respectively. The Y:S concentration was 180 g/1. Macronutrient compositions were calculated based on autolyzed yeast [MP Biomedicals, catalogue no. 103304 containing 62% protein]. Each solid diet contained 0.01% phosphoric acid and 0.1% proprionic acid as antimould agents and were prepared in sterile, distilled water.

#### Fly sampling

Newly-eclosed female flies were sorted and placed in longevity cages. Three replicate cages were run per diet (i.e., P:C 1:21, 1:11, 1:5, 1:2.5, 1:1, 3:1 and 1:0), each with 180 flies. Dead individuals were counted and removed from the cages every two days until all flies were dead. Life expectancy curves were analyzed using Log Rank Mantel-Cox tests. Ten live flies per treatment cage were sampled at 25%, 50% and 75% mortality (see Supplementary Fig. 1 and Supplementary Table 4). Flies were dissected (i.e, eggs removed), preserved in RNA later (Ambion) and stored at −80 °C for further analyses.

#### RNA extractions

We prepared up to three total RNA samples per dietary treatment by pooling individuals (n=10) of the same replicate cage. When less than 10 flies remained in the longevity cages, we discarded the sample. Subsequently, total RNA was extracted using a Trizol/RNeasy (Plus Mini kit, Qiagen) hybrid extraction protocol [see (Ponton et al., 2011b)]. Briefly, insects were homogenised in 1ml TRIzol reagent using a TissueLyser and 7mm stainless beads. Samples were incubated for 15min at room temperature and centrifuged for 10min at 12,000 × g at 4 °C. A standard volume of supernatant (800μl) was removed and added to 200μl of chloroform. Tubes were shaken vigorously for 15s, incubated at room temperature for 3min and centrifuged for 20min at 12,000 × g at 4°C. The aqueous phase (350μl) was transferred to a gDNA eliminator column from an RNeasy Plus Mini Kit (Qiagen) and all other steps were performed according to the manufacturer’s protocol (i.e. from step 4 in the version from Oct. 2005). Total RNA was eluted in 35μl of water. Extraction was followed by a DNase treatment (Ambion) to eliminate potential genomic DNA in the samples. RNA was then stored at −80°C before further processing. The quality and quantity of RNA was assessed with a Nanodrop ND-1000 spectrophotometer (Nanodrop Technologies). cDNA was produced using the QuantiTect Reverse Transcription Kit (Qiagen). cDNA was stored at -20°C until used.

#### Gene expression analysis

Gene expression was evaluated using custom made Taqman Low-Density Array (TLDA) Cards (Life Technologies/Applied Biosystems). Each TLDA card allowed for eight samples and assayed the expression of 21 immune genes (see Supplementary Table 4). Target gene expression levels were normalized using four reference genes (i.e., Ef1α100E, αTub84B, RpL32 and 18SrNA, see Supplementary Table 3). All samples were run on an ABI model 7900HT sequence detection system according to the protocol supplied by the manufacturer. Results were summarized using the 2^−ΔΔCt^ method. We log transformed the response variable before making statistical inferences, although all plots are of the raw data.

The effect of the percentage of dietary P:C was then tested for each gene and time point individually using generalized additive models (GAMs) that allowed for no a priori decision for choosing a particular response function. The percentage of protein in the diet was used as a descriptor of the diet composition.

### Statistical analyses

Statistical analyses were run using R (R Development Core Team, 2013) and SPSS (IBM Corp. released 2012. IBM SPSS Statistics for WINDOWS, v. 21.0. Armonk, NY: IBM Corp.).

## Author Contributions

FP and SJS designed the experiments. FP, KR and SSK ran the experiments. FP and JM analyzed the results. FP, JM, KR, SC, SSK, KW and SJS wrote the manuscript.

## Acknowledgments

The study was funded by the Australian Research Council (DP130103222 to SJS, KW and FP) and Macquarie University Research Seeding Grants (ID: 38277130 to FP).

---

**Supplementary Figure 1.**
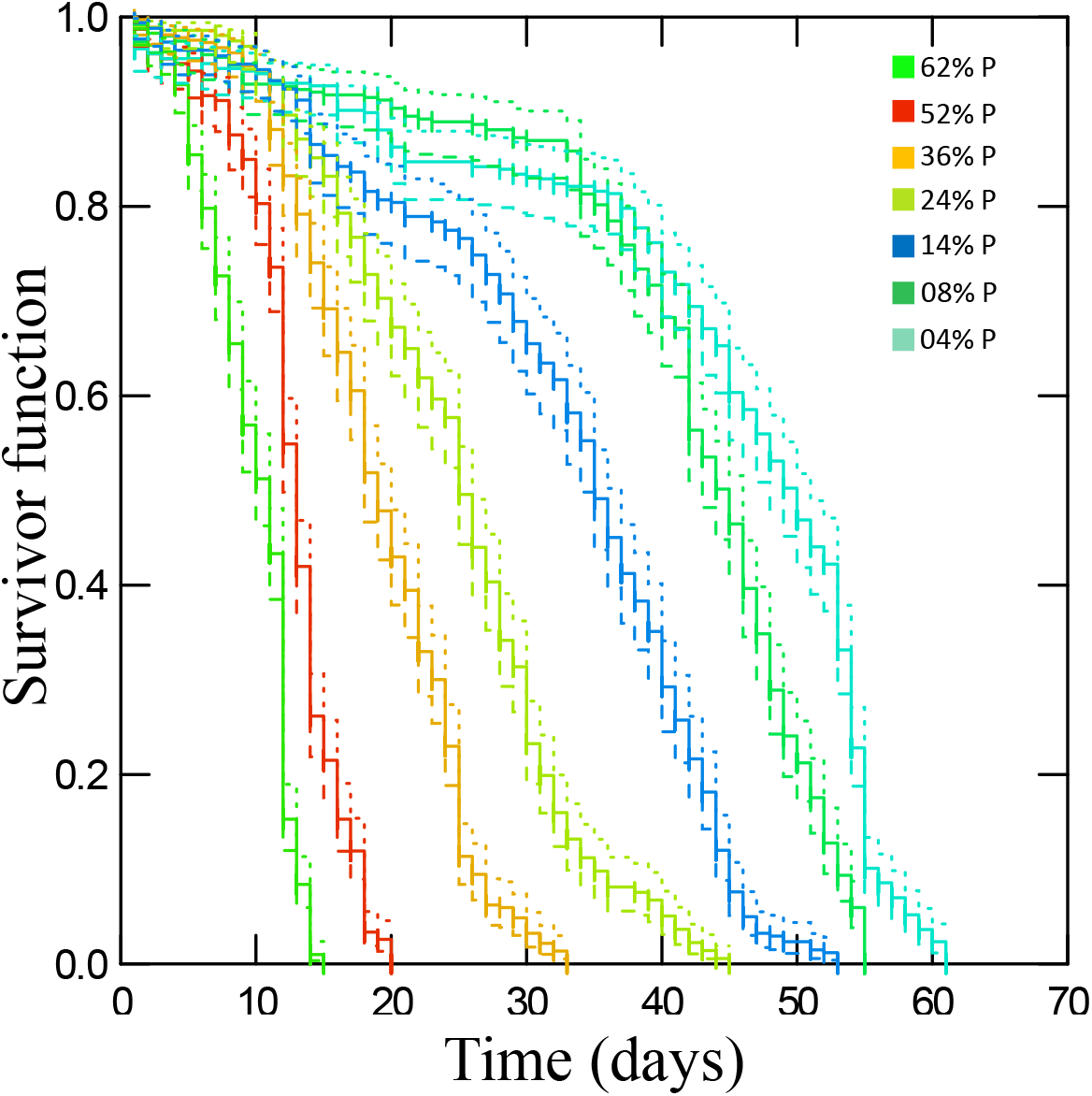
Life expectancy Kaplan–Meier curves with 95% confidence for flies fed seven diets varying in the percentage of protein (P) (Log Rank test, χ^2^=3166.5, df=6, p≤0.001).

**Supplementary Figure 2.**
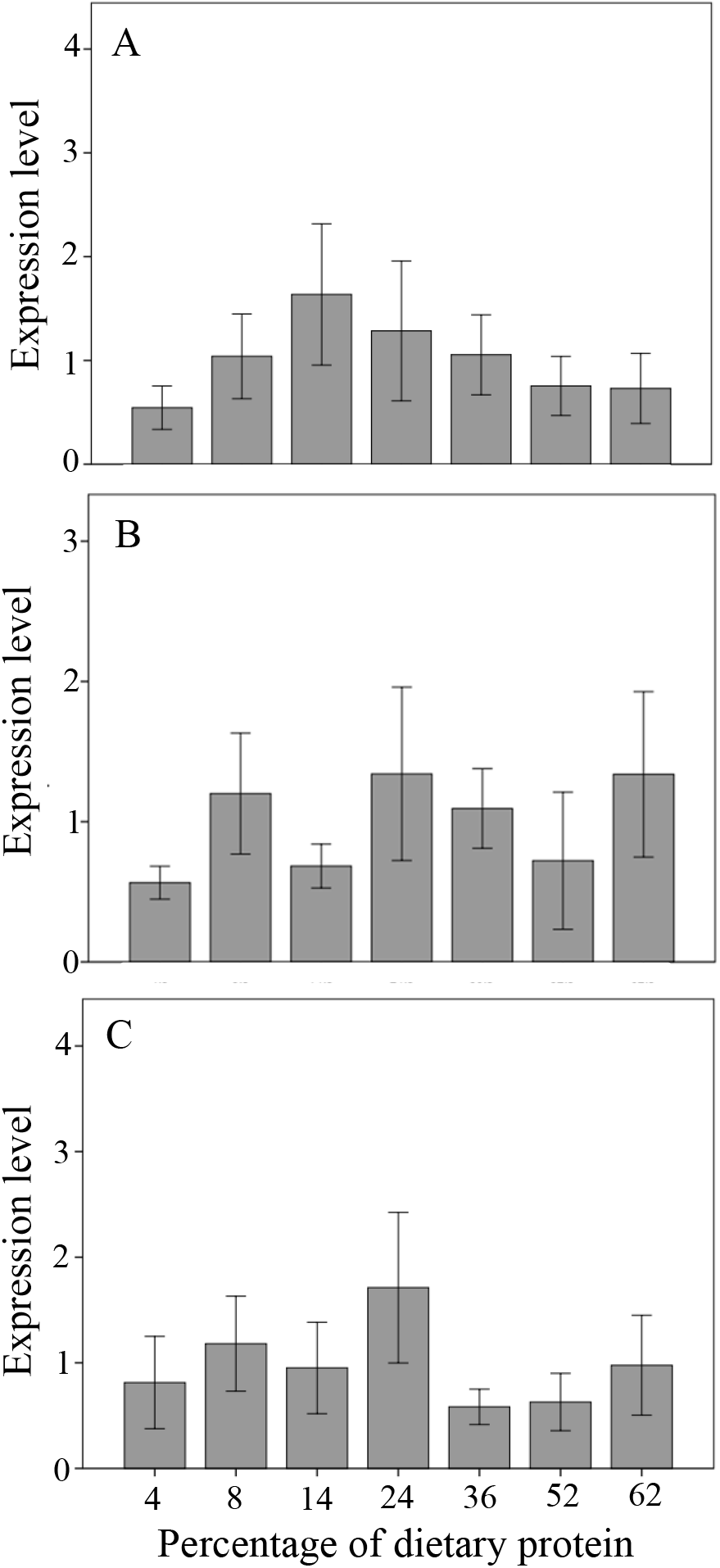
Expression levels of immune receptors (mean±SE) according to the percentage of protein in the diet at: A. 25% mortality, B. 50% mortality, C. 75% mortality.

**Supplementary Figure 3**. Estimated nonparametric smooths of immune gene expression levels from the generalized additive model according to the percentage of dietary protein at 25%, 50% and 75% mortality.

**Supplementary Figure 3 (1/4).**
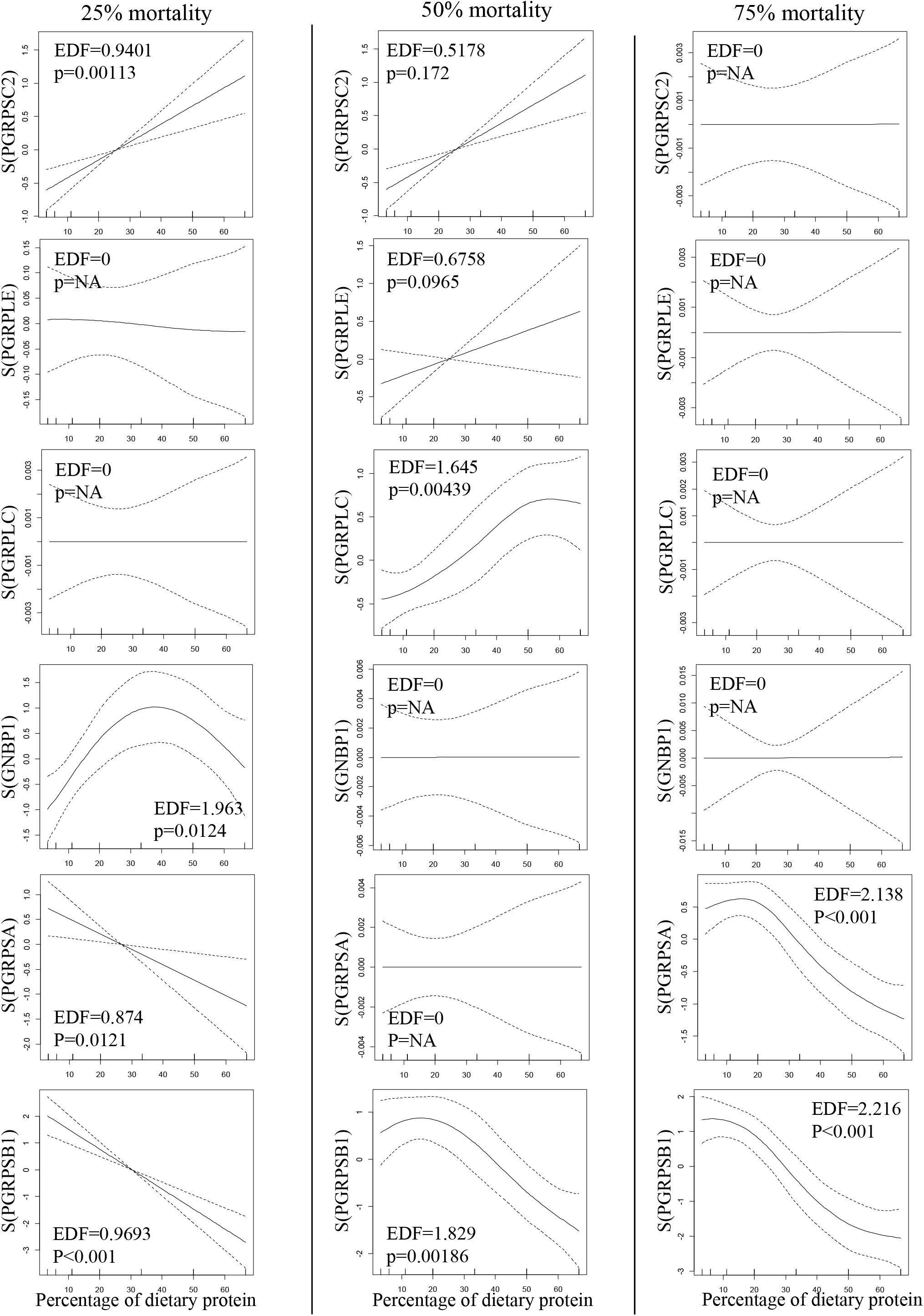
Molecules involved in the recognition of pathogens

**Supplementary Figure 3 (2/4).**
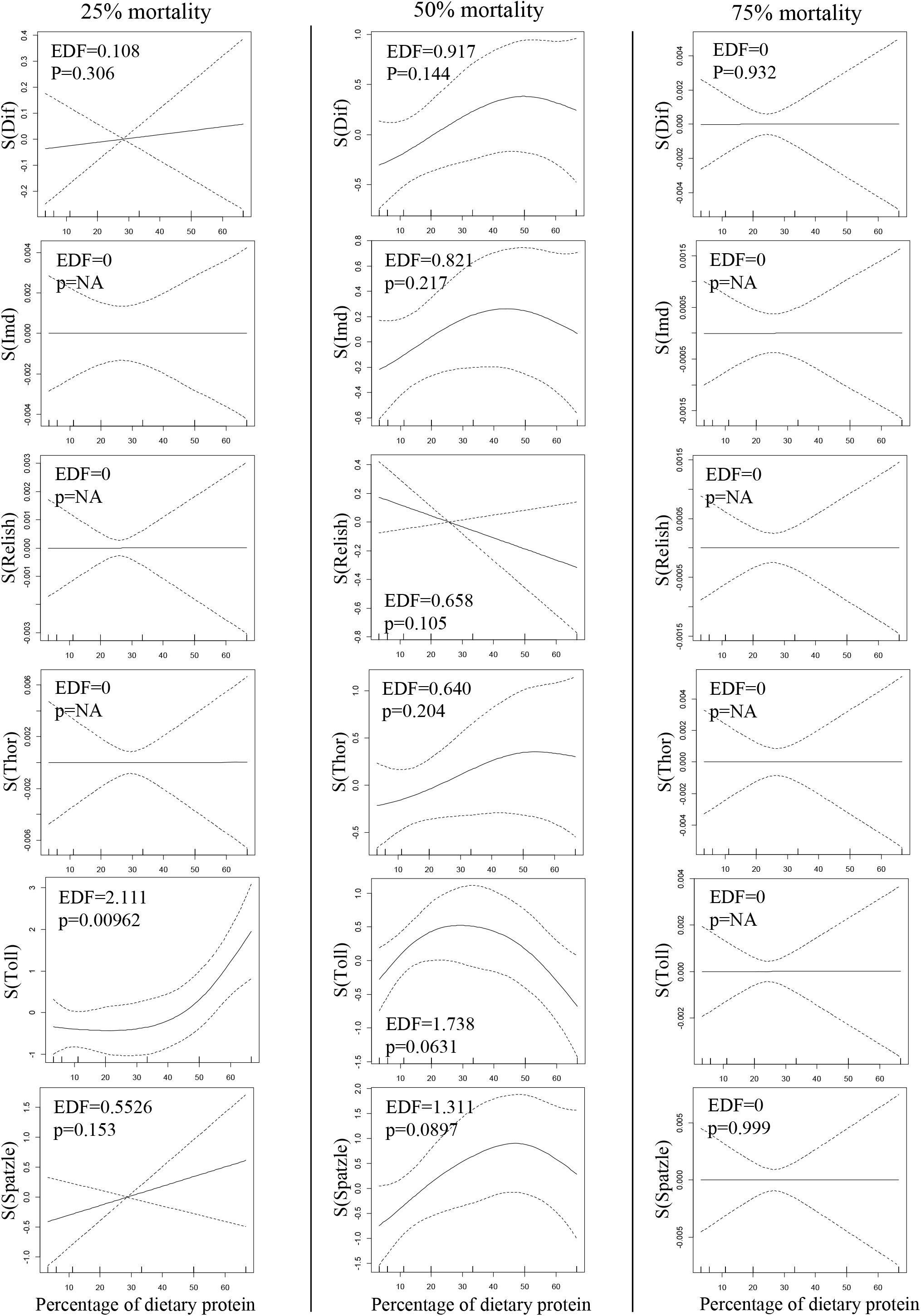
Molecules involved in the transduction of the immune signal

**Supplementary Figure 3 (3/4).**
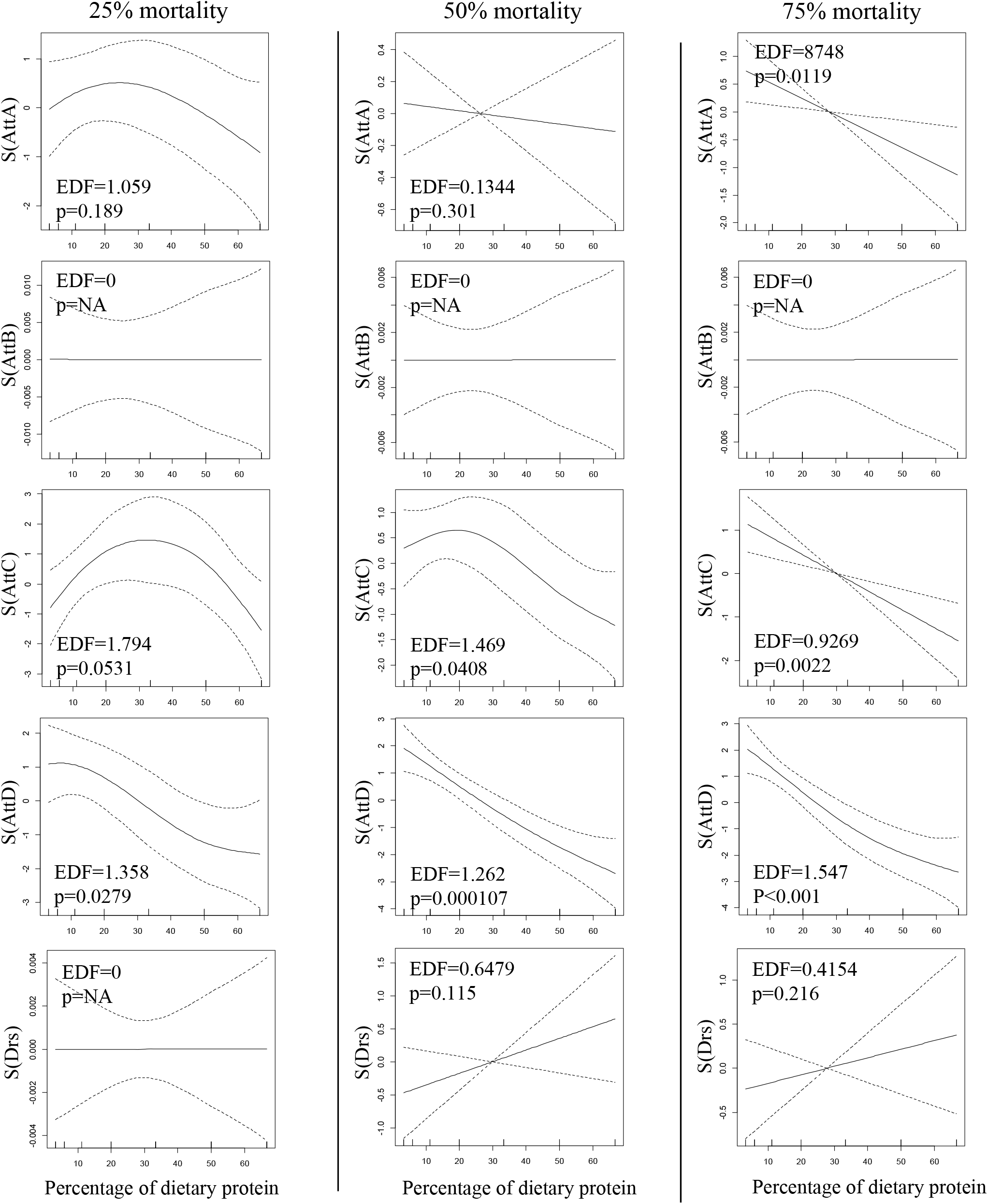
Antimicrobials

**Supplementary Figure 3 (4/4).**
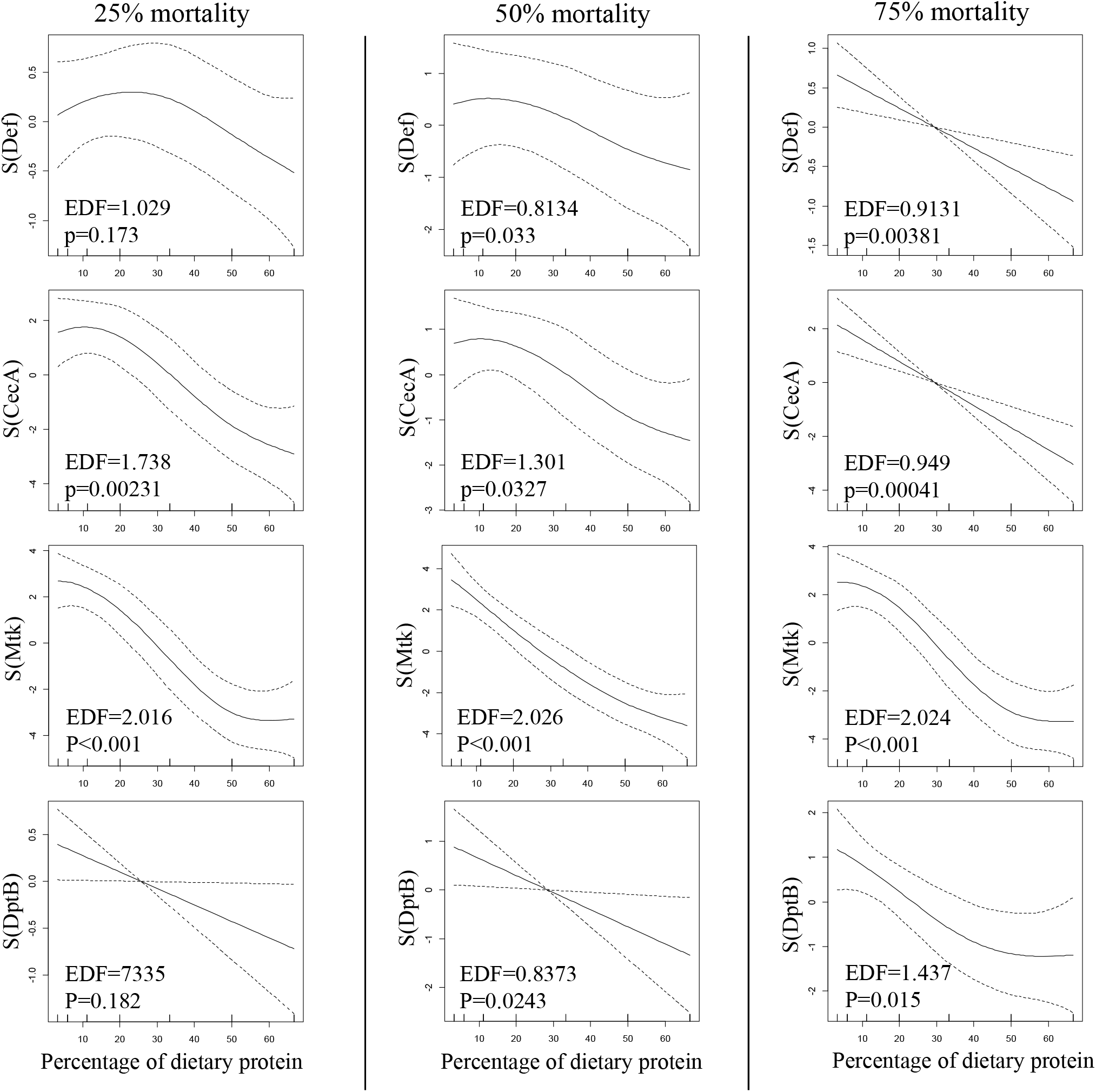
Antimicrobials

**Supplementary Table 1.** Macronutrient intakes for flies in the food choice experiment following three treatments (i.e., Control, Sham- and *M. luteus*-infected).

| Treatment | Total carbohydrate eaten (µg) | SD | Total protein eaten (µg) | SD | N |
| --- | --- | --- | --- | --- | --- |
| Naïve | 534.47 | 159.15 | 136.35 | 82.76 | 13 |
| *M. luteus* | 532.81 | 123.01 | 50.83 | 22.85 | 10 |
| Sham | 448.46 | 100.70 | 140.41 | 76.08 | 13 |

**Supplementary Table 2.**
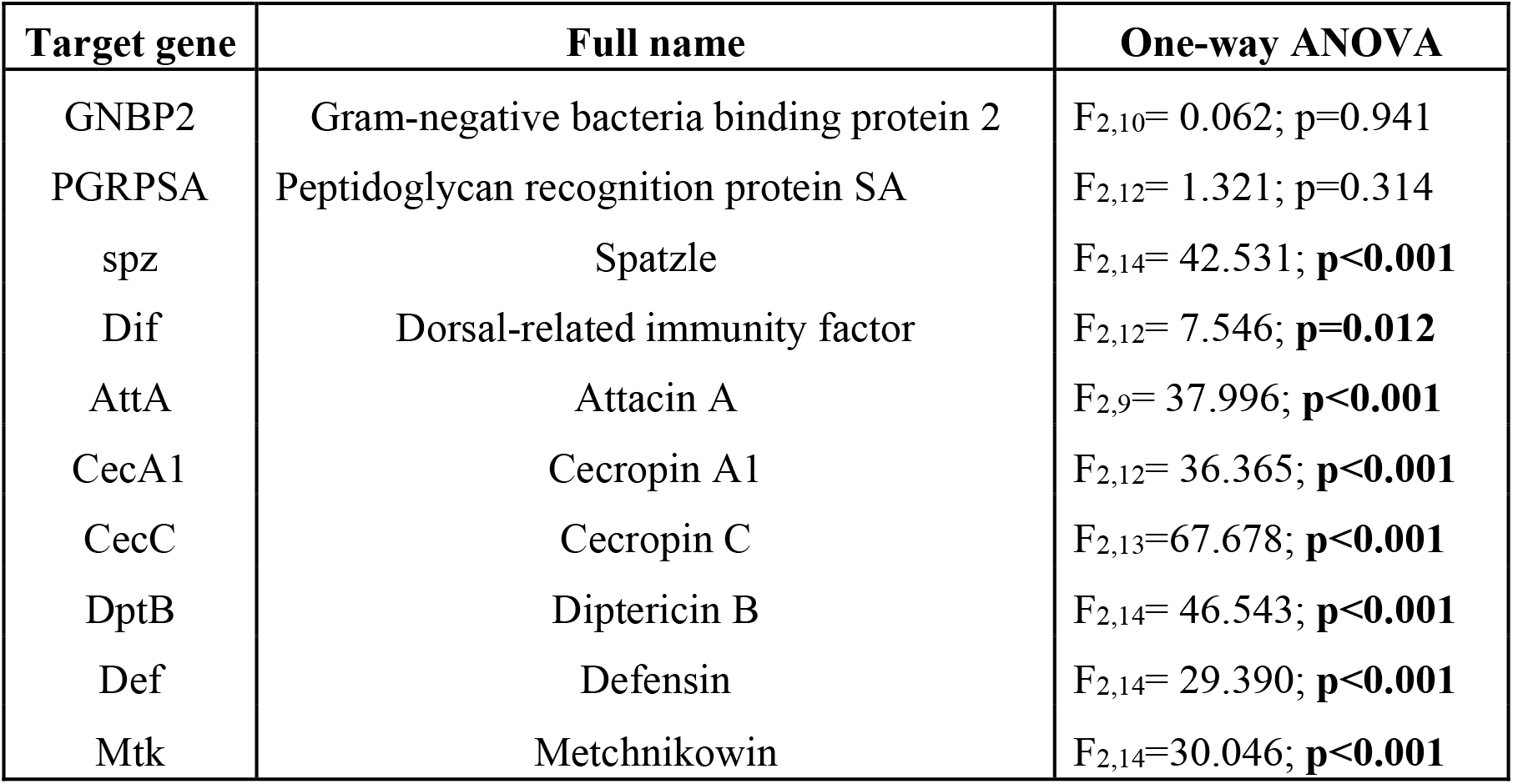
Gene details and statistical analyses following RT-qPCR assays.

**Supplementary Table 3.** **A**. Generalized linear model (Binomial error distribution) analyses to test for the effects of diet (low, medium and high P:C ratio) and treatment (*M. luteus*- and sham-infected, and naïve) on the number of dead flies after 15 days. **B**. Percentage of dead flies in each treatment and diet after 15 days.

**A**
| Factors | df | df residuals | Residuals deviance | p |
| --- | --- | --- | --- | --- |
| Diet | 2 | 1328 | 1260.2 | <0.001 |
| Treatment | 2 | 1326 | 1213.8 | <0.001 |
| Diet X Treatment | 4 | 1322 | 1191 | <0.001 |

**B**
| Diet | Treatment | Percentage of dead flies |
| --- | --- | --- |
| High P:C | Naïve | 97.04 |
| High P:C | Sham | 99.28 |
| High P:C | *M. luteus* | 94.56 |
| Medium P:C | Naïve | 21.79 |
| Medium P:C | Sham | 52.98 |
| Medium P:C | *M. luteus* | 62.59 |
| Low P:C | Naïve | 19.31 |
| Low P:C | Sham | 26.35 |
| Low P:C | *M. luteus* | 29.25 |

**Supplementary Table 4.**
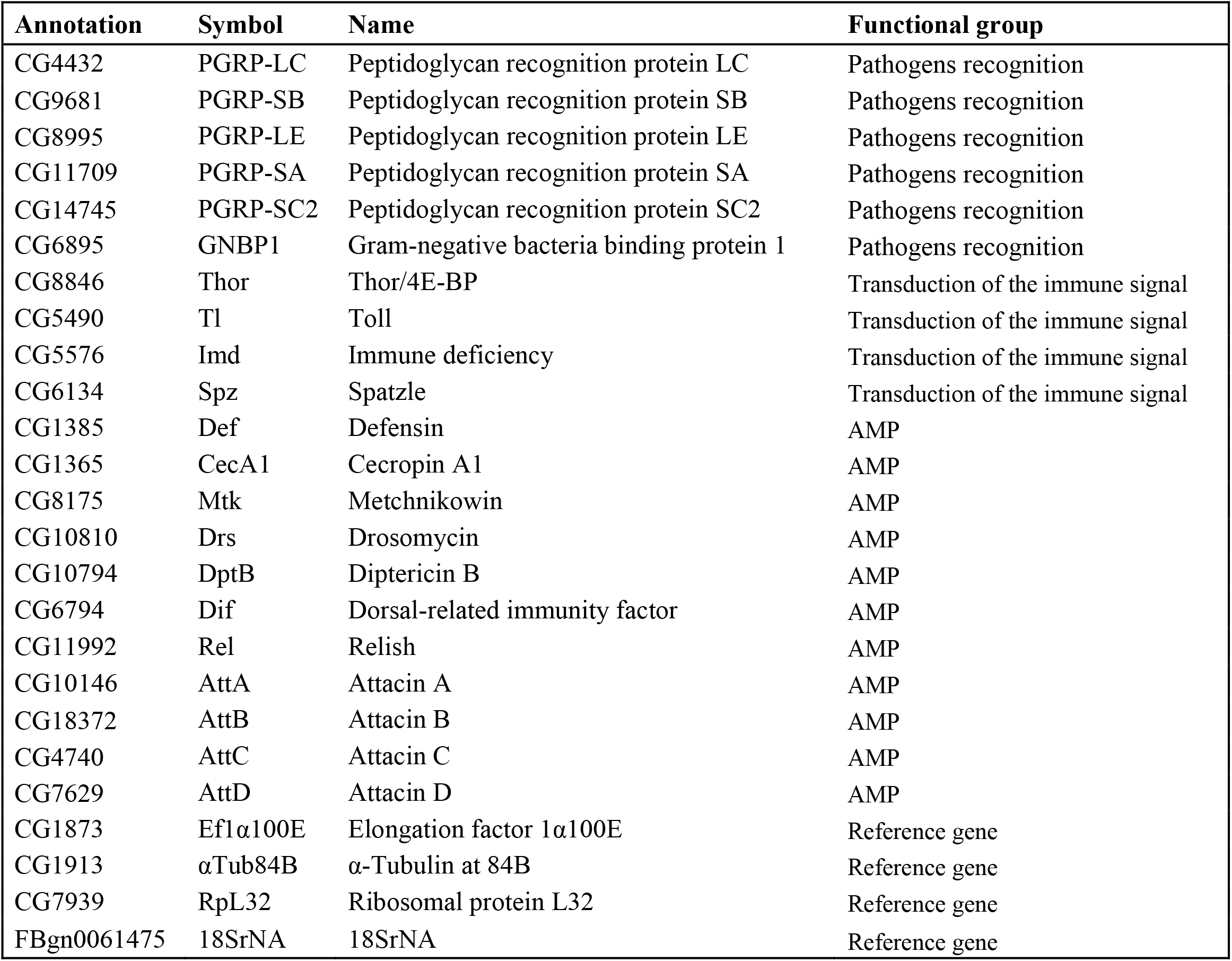
List of genes used in the design of the TLDA card assay.

**Supplementary Table 5.** Sampling at 25%, 50% and 75% mortality on the life expectancy curves for each of the seven diets varying in the protein-to-carbohydrate ratio.

| % Protein | 25% mortality |  | 50% mortality |  | 75% mortality |  |
| --- | --- | --- | --- | --- | --- | --- |
|  | average (days) | SD | average (days) | SD | average (days) | SD |
| 4 | 35.67 | 13.65 | 49.00 | 5.29 | 55.00 | 2.00 |
| 8 | 40.33 | 5.69 | 45.67 | 5.51 | 48.00 | 6.00 |
| 14 | 25.67 | 10.69 | 35.67 | 4.62 | 39.67 | 4.04 |
| 24 | 17.67 | 4.51 | 25.67 | 4.04 | 30.67 | 3.21 |
| 36 | 16.00 | 4.00 | 19.33 | 4.51 | 21.67 | 3.51 |
| 52 | 11.33 | 1.15 | 13.33 | 1.15 | 15.67 | 2.52 |
| 62 | 7.67 | 2.52 | 10.00 | 2.65 | 11.67 | 0.58 |

**Supplementary Table 6.** Kruskal-Wallis analyses to test for the effect of the percentage of dietary protein on the level of expression of immune receptors genes and genes coding for molecules involved in the transduction of the immune signal.

| Gene class | $\chi^2$ | df | *p* | N |
| --- | --- | --- | --- | --- |
| **25% mortality** |  |  |  |  |
| *Immune receptors* | 16.599 | 6 | **0.011** | 105 |
| *Transduction immune signal* | 12.653 | 6 | **0.049** | 126 |
| **50% mortality** |  |  |  |  |
| *Immune receptors* | 13.810 | 6 | **0.032** | 110 |
| *Transduction immune signal* | 11.097 | 6 | 0.085 | 134 |
| **75% mortality** |  |  |  |  |
| *Immune receptors* | 19.336 | 6 | **0.004** | 108 |
| *Transduction immune signal* | 5.944 | 6 | 0.429 | 125 |

**Supplementary Table 7.**
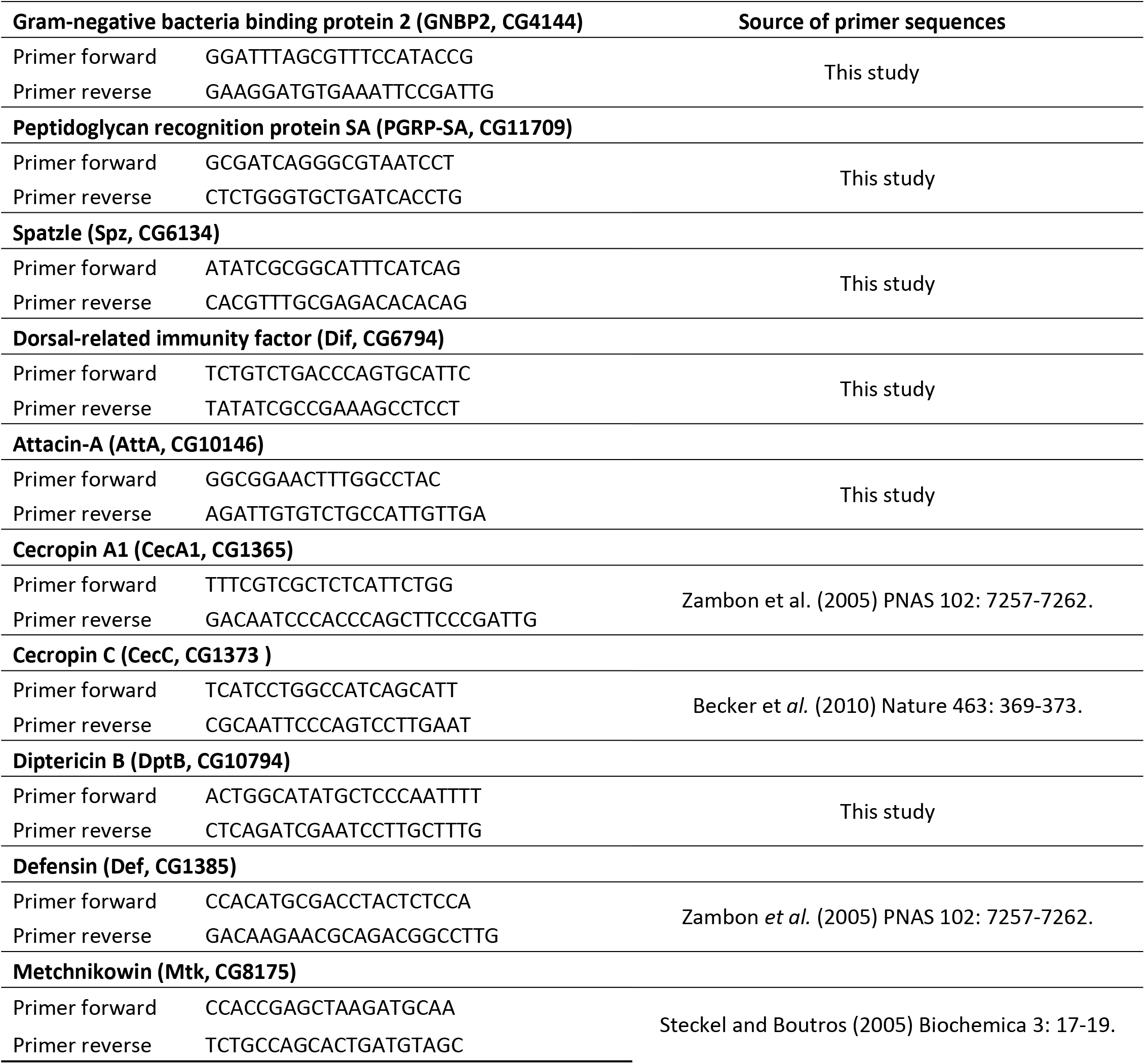
Primers of genes used in the RT-qPCR experiment.

